# Photosynthetic assimilate determines branch size and biomass more than branch number in Arabidopsis

**DOI:** 10.64898/2026.07.29.741629

**Authors:** Sungkyu Park, Scott A. Finlayson, Chenxin Li

## Abstract

Shoot branching is a primary determinant of plant form and crop yield, yet whether auxin or carbon supply is the proximal regulator of branching remains contested. In *Arabidopsis*, an earlier report that exogenous auxin fails to restore apical dominance after decapitation has been taken to weaken the case for auxin, and several studies have proposed that sugars are the primary regulator. Resolving this has been difficult because most perturbations of sugar status also disturb auxin. Here, we revisit the control of *Arabidopsis* branching using well-controlled, largely unperturbed plants and approaches designed to isolate each pathway. Contrary to the earlier report, apically applied auxin restored the suppression of rosette branching after decapitation, placing Arabidopsis in line with other species. In a dataset of 718 plants, cauline and rosette branching were weakly but significantly negatively correlated, consistent with a polar-auxin-transport-based model and contrary to a previous conclusion of no relationship. Removing all rosette leaves at bolting slowed bud growth but did not alter the final number of branches. Varying photosynthetic photon flux density across six natural accessions, analyzed by piecewise structural equation modeling, showed that photoassimilate acted far more strongly on the mass deposited into branches than on whether a bud initiates a branch. We conclude that auxin remains a major regulator of apical dominance in Arabidopsis, and that photosynthetic assimilate, while required as a substrate for branch growth, contributes little to determining branch number but more to branch size and biomass.

## Introduction

Shoot branching is a primary determinant of plant form, shaping how a plant displays its leaves, competes with its neighbors, and positions its reproductive structures. Branch number and position directly influence light capture, reproductive output, and harvest index. Branching has therefore been a major target of crop domestication and improvement, and much of the yield gain in cereals and horticultural species reflects changes in tillering or lateral shoot production (Mathan et al. 2016; Wang et al. 2018). Branching is also highly plastic, allowing plants to adjust their architecture in response to neighbor proximity, nutrient supply, defoliation, and other cues (de Jong et al. 2014; Kebrom and Mullet, 2015; Holalu et al. 2021). How this plasticity is achieved mechanistically remains actively debated after nearly a century of research.

Axillary buds form in the axils of leaves and either remain dormant or grow out to form branches. The classical framework accounting for their selective outgrowth, apical dominance, was established by Thimann and Skoog (1933), who showed that auxin produced at the shoot apex and transported basipetally through the polar auxin transport stream (PATS) suppresses the outgrowth of inferior buds. Later work refined this model and added further hormonal regulators, notably cytokinins (CK), which promote bud outgrowth, and strigolactones (SL), which suppress it (Domagalska and Leyser, 2011; Barbier et al. 2019). Two principal models describe how apically derived auxin acts at a distance. In the auxin canalization model, a bud grows out only once it establishes auxin export into the main stem, a capacity limited by the existing PATS (Prusinkiewicz et al. 2009; Balla et al. 2011). In the second-messenger model, stem auxin instead modulates the levels of CK and SL, which act directly on the bud (Brewer et al. 2009; Mason et al. 2014).

The standard physiological demonstration of apical dominance, that decapitation releases axillary buds and that replacing the apex with exogenous auxin restores their suppression, has been reproduced in numerous species (Thimann and Skoog, 1933; Cline, 1996). In *Arabidopsis*, however, exogenous auxin was reported not to restore apical dominance after decapitation (Cline, 1996), an apparent exception that has been taken to weaken the case for auxin in this species. Genetic and physiological evidence nonetheless implicates auxin in *Arabidopsis*: mutants compromised in auxin signaling show greatly reduced apical dominance (Stirnberg et al. 1999); auxin-overproducing lines branch less (Zhao et al. 2001); apically supplied auxin inhibits cauline bud growth in the split-plate assay (Chatfield et al. 2000); engineered rewiring of PIN1 responses to auxin suppresses branching (Khakhar et al. 2018); and manipulation of phyB/PIF signaling, which alters auxin responsiveness, strongly affects rosette bud activity (Reddy et al. 2014; Holalu et al. 2020, 2021). The tension between robust cross-species physiology and the equivocal direct demonstration in the principal model species remains unresolved.

Sugars have more recently been implicated as regulators of bud outgrowth. The early observation that the apex is a strong sink, and that bud release often coincides with greater sucrose availability, suggested that competition for carbon between apex and lateral buds could contribute to apical dominance (McIntyre, 1964). This idea has been elaborated by findings that decapitation rapidly raises sugar levels in axillary buds before measurable changes in auxin occur, and that exogenous sucrose releases buds from dormancy in pea, rose and sorghum, among other species (Kebrom et al. 2012; Mason et al. 2014; Barbier et al. 2015; Fichtner et al. 2017). These results have been read in two not-mutually-exclusive ways: a carbon-limitation model, in which sugars are required simply as a substrate for growth, and a sugar-signaling model, in which sugar status is an active regulatory input integrated with hormonal signals (Barbier et al. 2015, 2019; Otori et al. 2019; Patil et al. 2022; Beveridge et al. 2023).

A question that follows is whether the classical auxin model retains its explanatory power in light of these sugar-based models. Several reports have argued or implied that sugars, rather than auxin, are the primary or proximal regulator of bud outgrowth in Arabidopsis (Mason et al. 2014; Fichtner et al. 2021), though exogenous auxin was reported not to restore apical dominance in decapitated *Arabidopsis* (Cline, 1996), in apparent contrast to many other species. The two classes of model are not, however, inherently incompatible, and their relative contributions remain poorly resolved, in part because perturbations that alter sugar status frequently also perturb auxin transport, biosynthesis or signaling, and vice versa (Otori et al. 2017). Distinguishing them requires approaches that isolate, as far as possible, the contribution of each pathway and that probe the response of an essentially unperturbed plant.

Here, we revisit several basic questions about the control of *Arabidopsis* shoot branching to clarify the relative contributions of auxin and photosynthetic assimilate. We re-examine whether apical auxin can suppress bud outgrowth after decapitation, under conditions designed to address limitations of earlier work. We then ask whether rosette branching is correlated with cauline branching, as a PATS-based model predicts that cauline buds contribute to the auxin flux that suppresses inferior buds. Carbon supply is then assessed directly, by removing rosette leaves at bolting to reduce assimilate reserves and assimilation capacity, and by varying photosynthetic photon flux density (PPFD) across a range of natural accessions to test how far carbon availability dictates the number, length and biomass of branches. Together, our results indicate that auxin remains a major regulator of apical dominance in Arabidopsis, and that photosynthetic assimilate, while clearly required as a substrate for branch growth, contributes little to determining how many branches a plant produces but more to how much they elongate, and most to the material deposited into them.

## Results

### Rosette branching was negatively associated with cauline branch numbers

A prior study suggested cauline and rosette branching are not related (Fichtner et al. 2022). We re-examined this relationship in Col-0 using set of 718 records spanning 24 experiments, in which plants were grown under closely matched light, temperature, humidity, and medium conditions. Rosette leaf number varied substantially, rosette branch number varied modestly, and cauline branch number varied little (Fig. S1). Across the full dataset, cauline nodes/branches and rosette branch numbers were weakly but significantly negatively correlated (R² = 0.08, *p* = 1.27×10^-14^; Fig. 1a). To reduce the potential influence of flowering time, a core group of plants having the two most frequent number of rosette leaves, 10 and 11, was examined (n = 357). In this dataset, the negative correlation between cauline nodes/branches and rosette branches was stronger and still highly significant (R² = 0.16, *p* = 2.57×10^-15^; Fig. 1b). In the larger dataset, and also in all other Col-0 plants grown in our lab, all cauline nodes have produced branches without exception (Fig. 1c). Furthermore, of the 718 records in the dataset, the greatest number of plants produced 5 rosette branches, and every plant, except for 1, produced at least 3, in constrast to the consierably more variable number of rosette nodes formed (Fig. 1d,e). Thus, node position from the top, rather than stem morphology, is the better predictor of branch outgrowth, since virtually all of the upper 3 rosette nodes produce branches, just as all the cauline nodes do. This suggests that differences in bud fate between cauline and rosette reflects elevation in the plant rather than whether the node is associated with an elongated (cauline) or compressed (rosette) part of the stem.

**Figure 1.**
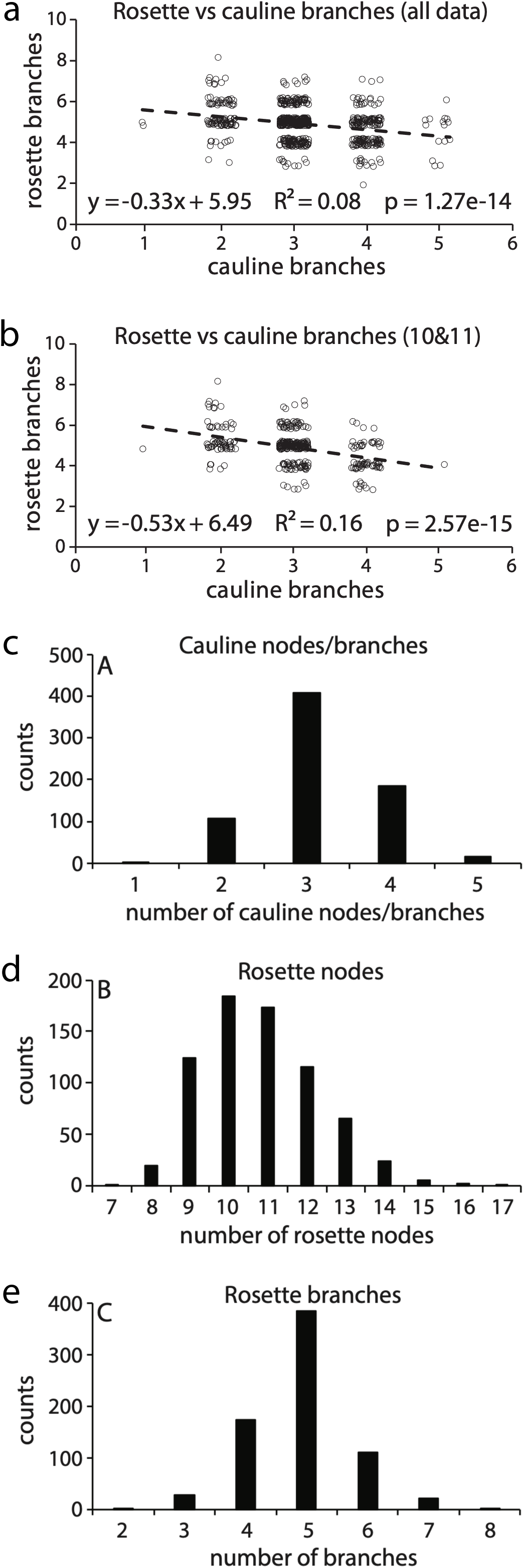
Rosette branching is negatively correlated with cauline branch numbers in Col-0. **(a)** Rosette branch numbers plotted against cauline node/branch number for all 718 records spanning 24 experiments (n = 718). **(b)** The same relationship for the core group of plants having the two most frequent rosette leaf numbers, 10 and 11 (n = 357). In (a) and (b), the dashed line is the least-squares regression, and the regression equation, coefficient of determination (R^2^), and *P*-value are given for each regression. **(c-e)** Frequency distribution of cauline node/branch number (c), rosette node number (d), and rosette branch number (e) across the full dataset. All cauline nodes produced branches, so cauline node and cauline branch numbers are equivalent. All plants were grown under very similar light, temperature, humidity, and medium conditions.

### Rosette branching was promoted by removing cauline branches

We next investigated whether the presence of cauline branch affects rosette branch formation. Removing the cauline branches promoted rosette branching from 5.3 branches in intact plants to 7.2 in plants with cauline branches removed (Fig. 2a,b). Removing the cauline branches also promoted elongation of the rosette branches that has initiated, and this effect was evident from the top to the bottom of the rosette (Fig. 2c,d). It may be concluded that cauline branches inhibit the initiation of branches at lower rosette positions and suppress rosette branch elongation throughout the rosette. These conclusions agree with many prior reports of correlative inhibition across variety of species, but do not define the underlying mechanism. The data are consistent with both branch regulation via the PATS and by sugar availability, two currently popular hypotheses employed to describe branch patterning.

**Figure 2.**
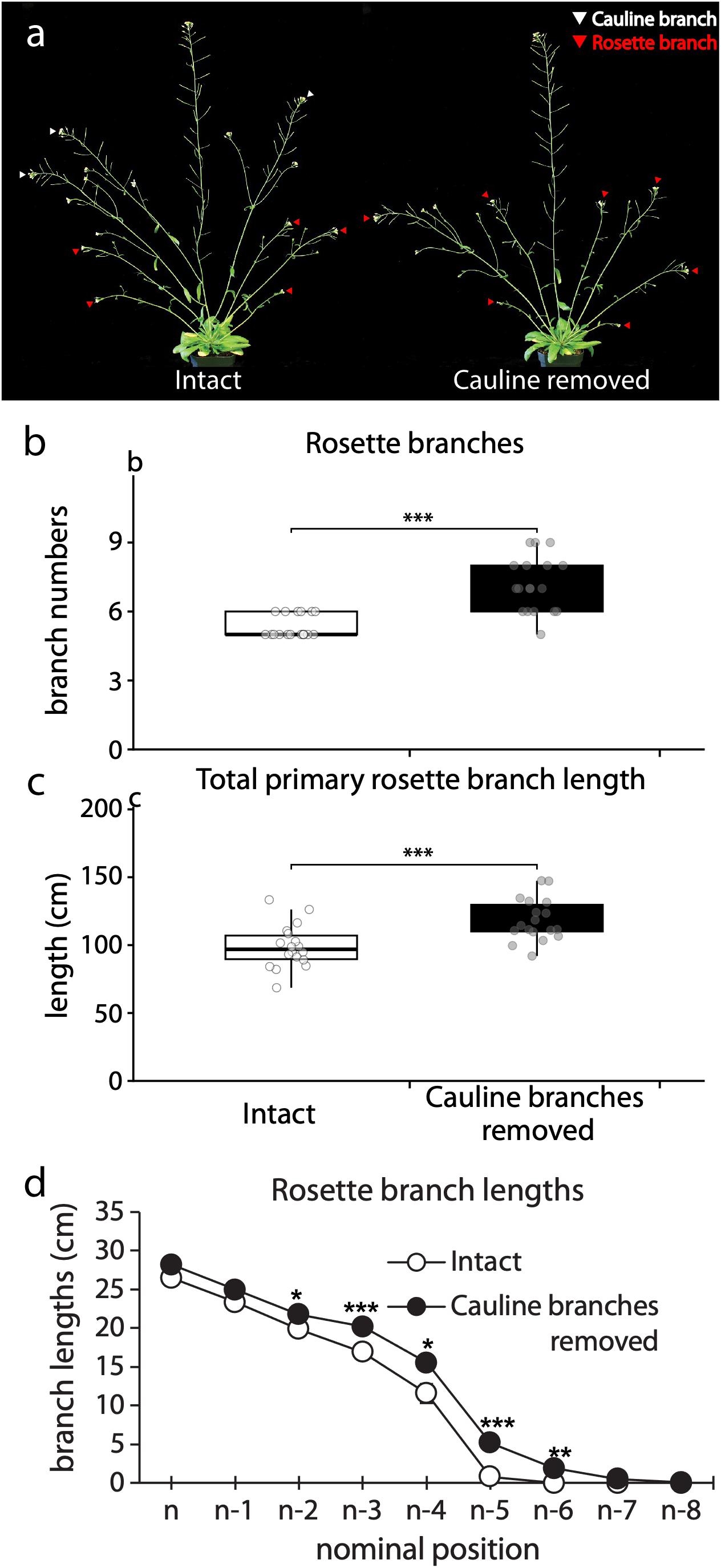
Removal of cauline branches promotes rosette branch initiation and elongation. **(a)** Representative Col-0 plants showing shoot architecture at 10 days after anthesis, either intact (left) or with all cauline buds removed at anthesis (right). **(b)** Rosette branch number and **(c)** total primary rosette branch length in intact plants and plants with cauline branches removed. **(d)** Rosette branch lengths by bud position, where position n is the uppermost rosette node and n-8 the lowest. n = 18 plants per treatment. Asterisks indicate differences between treatments by Student’s *t*-test: * *P* < 0.05, ** *P* < 0.01, *** *P* < 0.001.

### Apically applied auxin suppressed decapitation-induced rosette branching

Under the PATS hypothesis, auxin transported from the apex would be expected to mediate this inhibition. A prior study found that apically applied auxin did not restore apical dominance in decapitated *Arabidopsis*, raising questions about the role of auxin in this system (Cline, 1996; Cline et al. 2001). We tested this by decapitating plants shortly after bolting and applying the synthetic auxin NAA to the cut stem (Fig. 3a). Rosette branch numbers increased from 5 in intact plants to 7.9 in decapitated plants, and NAA applied to the cut stem restored branching to intact levels (5.3; Fig. 3b). Furthermore, decapitation combined with NAA decreased the elongation of the buds that grew to form branches (Fig. 3c,d). Thus, as has been shown in many other species, supplying an apical auxin source re-established the suppression of rosette branching in decapitated *Arabidopsis*.

**Figure 3.**
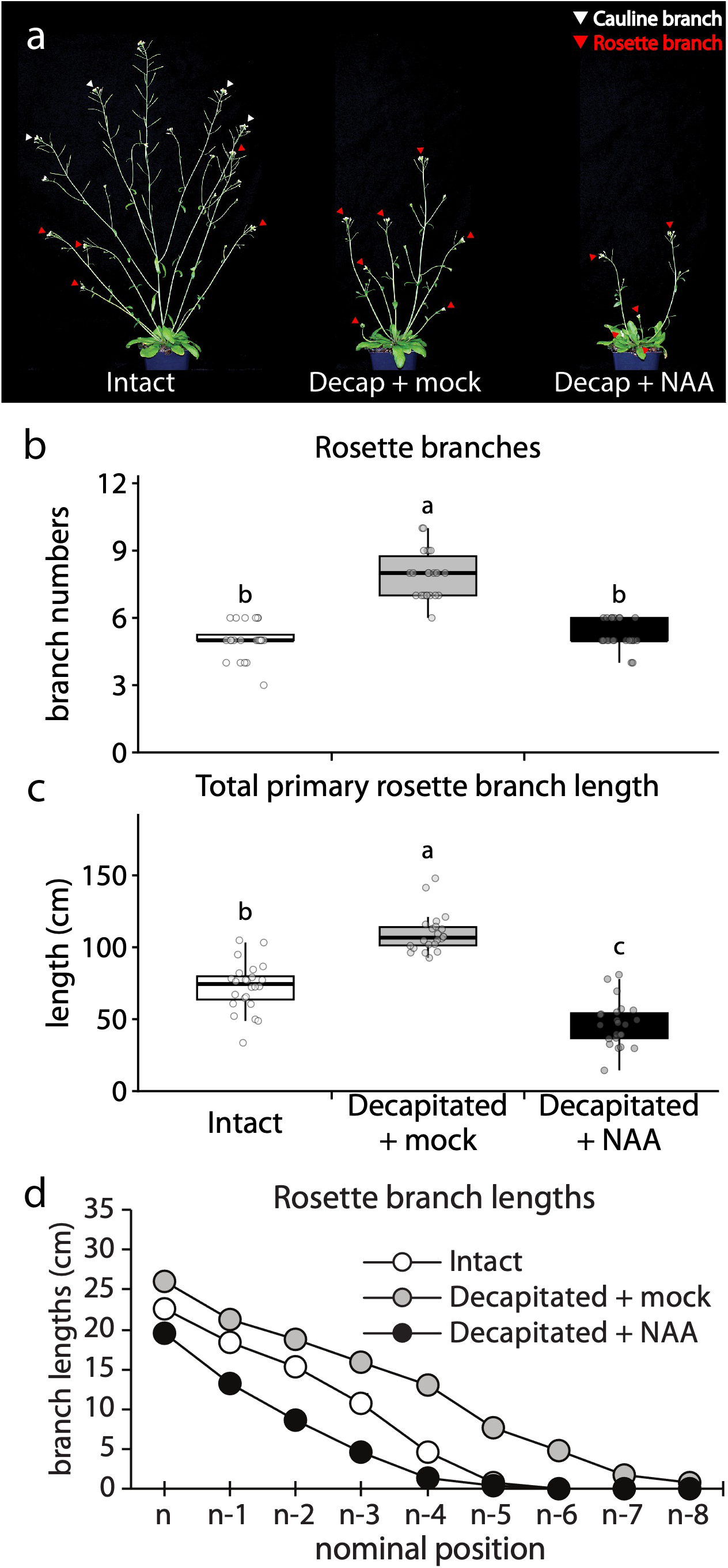
Apically applied auxin suppresses decapitation-induced rosette branching. **(a)** Representative Col-0 plants showing shoot architecture at 10 days after anthesis: intact (left), decapitated at anthesis (middle), or decapitated at anthesis with NAA applied to the cut stem (right). **(b)** Rosette branch number and **(c)** total primary rosette branch length in intact plants (white), decapitated plants (grey), and decapitated plants treated with NAA (black). **(d)** Rosette branch lengths by bud position, where position n is the uppermost rosette node and n-8 the lowest. n = 22-24 plants per treatment. Different letters indicate significant differences between treatments (Tukey’s HSD, *P* < 0.05).

### Photosynthetic carbon assimilation is weakly associated with branch initiation in Col-0

Our data are consistent with a major role for auxin in regulating *Arabidopsis* branching. However, the potential contribution of sugars produced by the photosynthetic carbon reduction cycle to branching remains obscure. Evidence from diverse species and experimental approaches supports a role for sugar availability in regulating bud outgrowth. Yet the effect of photosynthetic assimilate production on branching has not been directly tested in *Arabidopsis* by varying Photosynthetic Photon Flux Density (PPFD), an approach that avoids the confounding effects of exogenous sugar application and the growth inhibition associated with sugar deprivation.

We examined the role of sugars produced via the photosynthetic carbon reduction cycle in Col-0 branching. Neutral density filters permitted the provision of varying levels of PPFD without altering other environmental parameters, such as light spectrum or photoperiod, that might exert additional effects on the plants. The initial experiment provided low (120 μMoles m^-2^ s^-1^), medium (180) and high (240) PPFD from 20 days after sowing (DAS), the onset of bolting (Fig. S2d). It was anticipated that increased photosynthesis just prior to bud formation would stimulate branching, including branch initiation, without dramatically altering other aspects of development. Higher PPFD significantly increased shoot DW, but had little effect on branch initiation (Fig. S2). The experiment was repeated with light treatments applied from 6 DAS rather than from bolting at 20 DAS (Fig. 4a). Elevated light had only a slight impact on branch initiation but a stronger positive effect on total branch elongation (Fig. 4b,c). Higher resolution assimilate partitioning was obtained in this experiment by measuring rosette leaf, main shoot and branch/bud DW separately. Higher PPFD increased the DW commensurately (Fig. 4d). The specific masses (μg DW mm^-1^) of the longest branch and the main stem were also increased with PPFD treatment (Fig. 4e,f).

**Figure 4.**
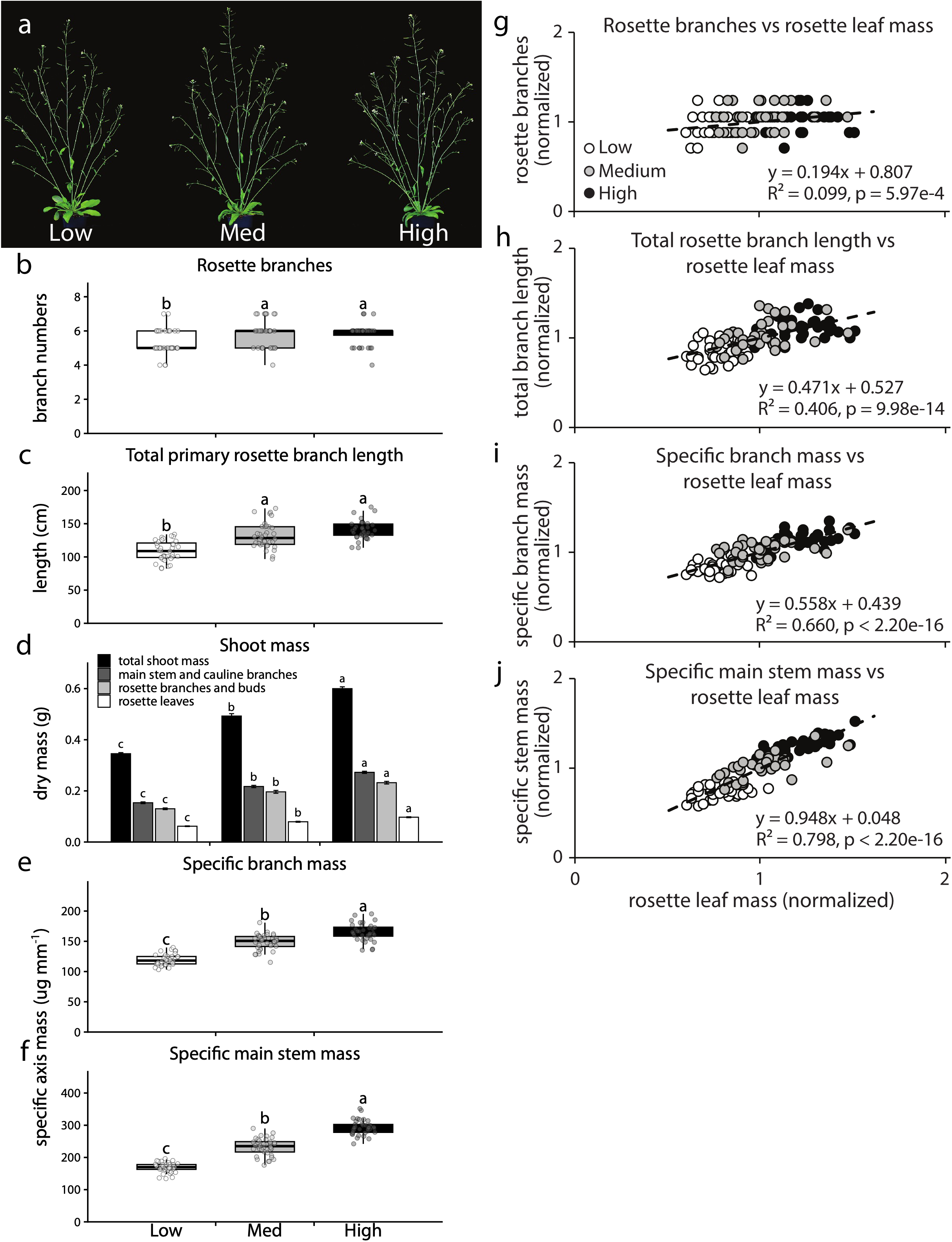
Photosynthetic assimilation is weakly associated with rosette branch initiation but strongly associated with branch elongation and material deposition in Col-0. Plants are grown at low (120 µMoles m^-2^ s^-1^), medium (180 µMoles m^-2^ s^-1^), or high (240 µMoles m^-2^ s^-1^) PPFD from 6 days after sowing and harvested 10 days after anthesis. **(a)** Representative Col-0 plants showing shoot architecture under varying PPFD treatment. **(b)** Rosette branch number. **(c)** Total primary rosette branch length. **(d)** Shoot dry mass partitioning among total shoot, main stem and cauline branches, rosette branches and buds, and rosette leaves. **(e)** Specific branch mass. **(f)** Specific main stem mass. **(g-j)** Regressions of rosette branch number (g), total rosette branch length (h), specific branch mass (i), and specific main stem mass (j) against rosette leaf mass, used as a proxy for photosynthetic assimilation. In (g)-(j), each parameter was normalized to its sample mean, which shows the relationships on a per-plant basis without altering the coefficient of determination (R^2^) or the *p*-value. Each point represents a single plant, shaded by PPFD treatment: low (white), medium (grey), or high (black). Dashed lines are least-squares regressions, and the regression equation, coefficient of determination (R²), and *P*-value are given for each regression. n = 36 plants per treatment. Different letters indicate significant differences between treatments (Tukey’s HSD, *P* < 0.05). In (d), data are means ± SE, and treatments are compared separately within each mass fraction.

For each record, branch initiation (numbers) was regressed against rosette leaf DW. Rosette leaf DW reflected the shoot photosynthetic assimilation during the vegetative phase, the reservoir of accumulated carbon or sugars, and photosynthetic potential for future development. Branching parameters generally showed higher correlations with rosette leaf DW than with total shoot mass or main shoot mass (Fig. 4g,h; Fig. S2f,g). Weak, but significant correlation was observed between branch numbers and rosette leaf DW (Fig. 4g). Normalizing each parameter to its sample mean reveals the relationships on a per-plant basis without altering the coefficient of determination or the *p*-value. Other parameters, including total primary rosette branch lengths, specific branch mass, and specific main shoot mass, showed much greater correlations with, and responses to, rosette leaf mass (Fig. 4h-j). Overall, photosynthetic assimilation was only weakly associated with rosette branch initiation but more strongly associated with branch lengths and the amount of material deposited into the axes (buds and branches) formed.

### Architectural responses to photosynthetic carbon assimilation in select *Arabidopsis* accessions: increased PPFD uniformly promoted material deposition into axes, but had varying effects on branch initiation

The experiment was revised and standardized, with varying light treatments imposed uniformly at 10 DAS, to assess the responses of a small selection of *Arabidopsis* accessions including Col- 0, Ob-0, Chi-0, Van-0, Bay-0 and Sha. These accessions were selected to expand the geographic and climatic range of *A. thaliana*, allowing to test whether the role of carbon availability in branching regulation generalizes across the species’ natural variation. Overall, little difference in shoot architecture was apparent with varying PPFD within accessions (Fig. 5a).

**Figure 5.**
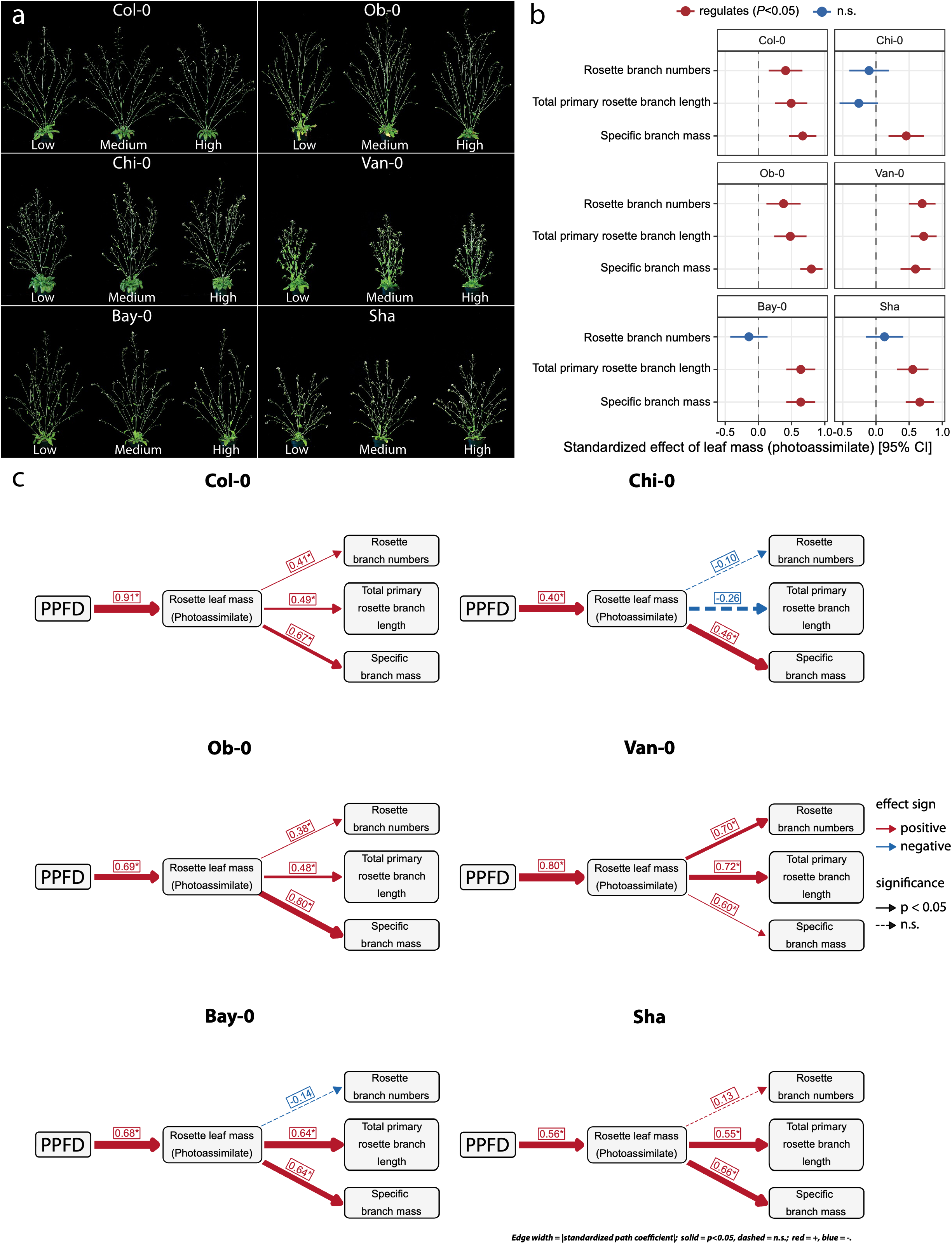
Photoassimilate acts consistently on branch mass and elongation but variably on branch initiation across *Arabidopsis* accessions. Six accessions (Col-0, Ob-0, Chi-0, Van-0, Bay-0, and Sha) are grown at low (120 µMoles m^-2^ s^-1^), medium (180 µMoles m^-2^ s^-1^), or high (240 µMoles m^-2^ s^-1^) PPFD from 10 days after sowing and harvested 10 days after anthesis. **(a)** Representative plants of each accession under the three PPFD treatments. **(b)** Standardized effect of rosette leaf mass, used as a proxy for photoassimilate, on rosette branch number, total primary rosette branch length, and specific branch mass, shown for each accession. Points are standardized path coefficients and horizontal lines the 95% confidence intervals. Red denotes a significant effect (*P* < 0.05) and blue a non-significant effect. The dashed vertical line marks zero. **(c)** Path analysis of the PPFD → rosette leaf mass → branching architecture pathway, fitted separately for each accession by piecewise structural equation modeling. Boxes represent measured variables, and arrows represent unidirectional relationships among variables. Red arrows denote positive and blue arrows negatives relationships. Solid arrows denote significant paths (*P* < 0.05) and dashed arrows non-significant paths. Arrow thickness is scaled to the magnitude of the standardized regression coe cient, given beside each arrow. n = 51-54 plants per accession (n = 17-18 per PPFD treatment).

To integrate the responses across accessions, the PPFD → rosette leaf mass → branching architecture pathway was modeled by path analysis, with rosette leaf mass as a proxy for photoassimilate (Fig. 5b,c; Lefcheck, 2016). PPFD had a strong positive effect on rosette leaf mass in every accession (standardized regression coefficients 0.40-0.91; Fig. 5c), confirming that the light treatments altered assimilate supply as intended. The downstream effect of leaf mass, however, depended on the branching parameters. Leaf mass was consistently associated with the two mass-related parameters: specific branch mass in all six accessions (0.46-0.80) and total primary rosette branch length in five (0.48-0.72; non-significant in Chi-0). Its association with rosette branch numbers was weaker and present in only three accessions (Col-0, Ob-0, and Van- 0; 0.38-0.70), and absent in Chi-0, Bay-0, and Sha (Fig. 5b). With the exception of Van-0, where leaf mass affected branch number as strongly as it affected branch mass and length, the effect of photoassimilate on branch initiation was consistently smaller than its effect on material deposition into existing axes. This pattern suggests that carbon supply acts more on how much mass is deposited into branches than on whether a bud initiates a branch, consistent with a role for assimilate as a substrate for branch growth rather than as a primary determinant of branch number.

Rosette branch numbers increased slightly with PPFD in Col-0 and Van-0, while the other accessions were largely unresponsive (Fig. S3). Regression analysis confirmed a weak association between branch initiation and rosette leaf mass in Col-0 and a similar response in Ob-0 (Fig. S3g,h). No association was detected in Chi-0, while Van-0 showed a relatively strong association (Fig. S3i,j). In the case of Bay-0 and Sha, rosette branch numbers correlated more strongly with rosette leaf number than with rosette leaf mass (Fig. S4a-d). To control for the dominant association between leaf numbers and branching, branch initiation in these accessions was assessed as branches axil^-1^. On this basis, there was no correlation between rosette leaf mass and rosette branches axil^-1^ in both Bay-0 and Sha (Fig. S4e,f).

Varying PPFD had a modest effect on the total primary rosette branch lengths of Col-0, Van-0 and Sha (Fig. S5a-f). Regressions provided modest positive associations between rosette leaf mass and total primary rosette branch lengths for Col-0, Ob-0, Bay-0, and Sha (Fig. S5g-l). Van- 0 showed a strong positive association, while the association in Chi-0 was weak and negative.

Specific branch mass increased with PPFD in all accessions, although mean separation was incomplete in Col-0 and Chi-0 (Fig. S6a-f). Moderate to strong positive correlations between rosette leaf mass and specific branch mass were found for all accessions with the exception of Chi-0 where the relationship was still significant, but weaker (Fig. S6g-l). Specific main stem mass of all accessions showed a positive response to increasing PPFD except Chi-0 (Fig. S7a-f), and correlated strongly with rosette leaf mass in every accession (Fig. S7g-l).

### Defoliation of rosette leaves did not affect rosette branch initiation of Col-0

The data above suggested that bud outgrowth in Col-0 is largely determined by developmental programming and was only weakly responsive to carbon status. This possibility was investigated more aggressively by targeted dissection of the Col-0 shoot at the onset of bolting (Fig. 6a) and assessing architecture 12 days later (Fig. 6b). As expected, removing the main inflorescence (decapitation) increased the number of branches produced (Fig. 6c). After removal of all rosette leaves (defoliation) and therefore the major source of carbon, bud outgrowth appeared to arrest for about 3 days during which time the bud leaves expanded and presumably became photoautotrophic. At the termination of the experiment, intact and defoliated plants had produced equivalent numbers of branches (Fig. 6c). The combination of decapitation and defoliation resulted in a similar number of branches as decapitation alone, indicating that rosette leaf-derived carbon is dispensable for branch initiation but main shoot-derived signals exert strong effects.

**Figure 6.**
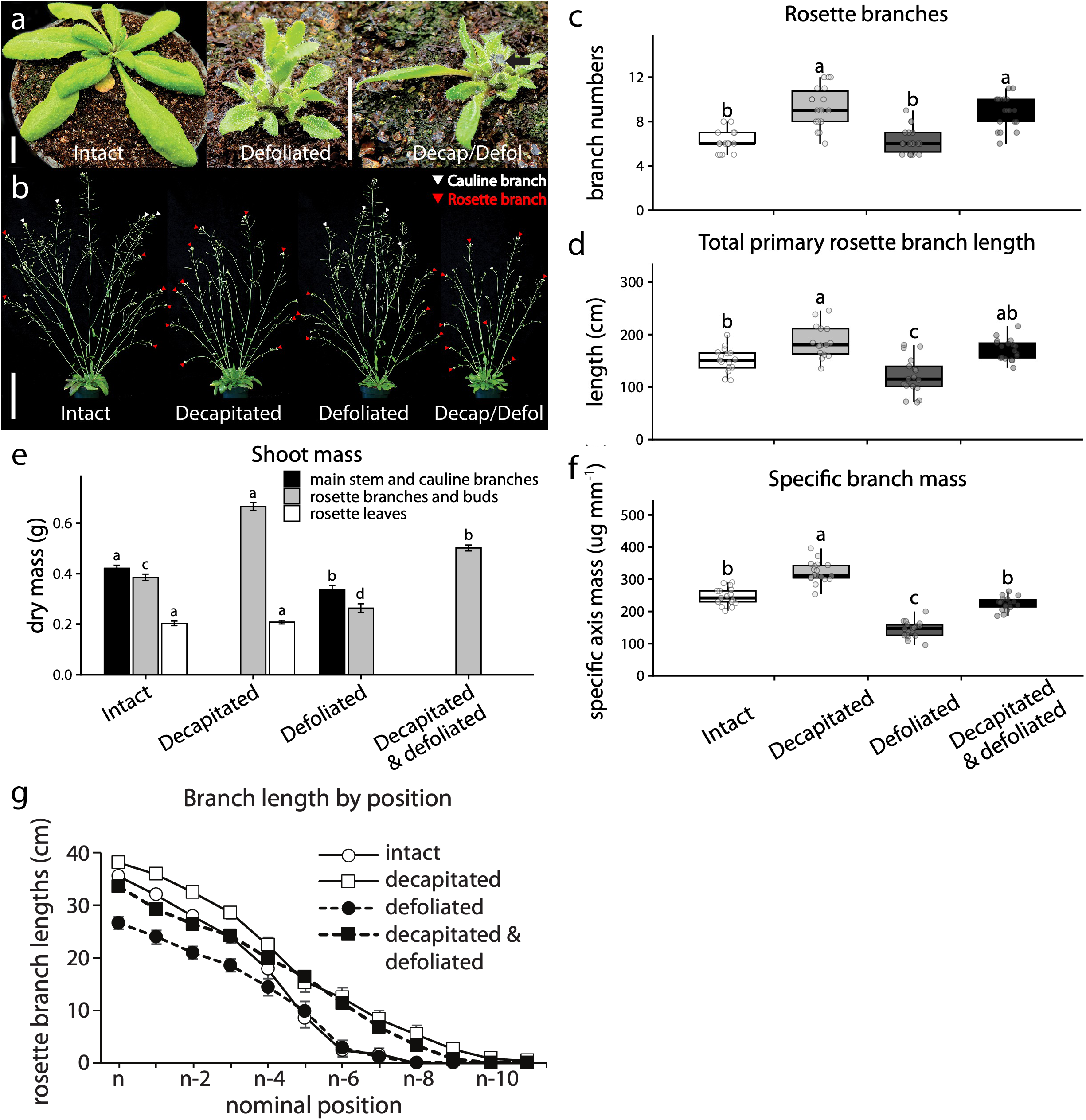
Defoliation did not alter rosette branch numbers in Col-0. Plants are treated with when the main stem began to bolt and assessed 12 days later. **(a)** Representative Col-0 plants immediately after treatment: intact (left), defoliated (all rosette leaves removed), and decapitated and defoliated. The black arrow marks the cut main stem. **(b)** Representative Col-0 plants showing shoot architecture at harvest. White arrowheads mark cauline branches and red arrowheads rosette branches. Scale bars = 1 cm in (a) and 10 cm in (b). **(c)** Rosette branch number. **(d)** Total primary rosette branch length. **(e)** Shoot dry mass partitioning among total shoot, main stem and cauline branches, rosette branches and buds, and rosette leaves. **(f)** Specific branch mass. **(g)** Rosette branch lengths by bud position, where position n is the uppermost rosette node and n-10 the lowest: intact (white circles, solid line), decapitated (white squares, solid line), defoliated (black circles, dashed line), and decapitated and defoliated (black squares, dashed line). In (e) and (g), data are means ± SE. Different letters indicate significant differences between treatments (Tukey’s HSD, *P* < 0.05). In (e), treatments are compared separately within each mass fraction. Fractions removed by a treatment are necessarily absent: the main stem and cauline branches from decapitated plants, and the rosette leaves from defoliated plants. n = 18 plants per treatment.

Decapitation also increased the total length of the rosette branches produced, the total DW of the branches, and the specific branch mass (Fig. 6d-f). Defoliation resulted in opposite responses, with the greatest reduction observed in the total branch length and specific branch mass. The combination of decapitation and defoliation slightly reduced elongation of the upper branches and promoted elongation of the lower branches compared to intact, producing a numerically, but not significantly, greater total branch length (Fig. 6d,g). However, the combined decapitation and defoliation treatment did significantly promote branch DW even though specific branch mass was not different from the intact treatment (Fig. 6e,f).

## Discussion

The classical view of branching regulation, apical dominance, proposes that auxin in the PATS inhibits the growth of inferior buds (Thimann and Skoog, 1933). This model has been modified and updated with new mechanistic interpretations and roles for additional hormones, but its core remains an auxin-dominated mechanism. This is not the only model for apical dominance however, and more recently models have been described whereby sugar availability and/or sugar signaling determines the degree of bud growth or inhibition (McIntyre, 1964; Kebrom et al. 2012; Mason et al. 2014; Barbier et al. 2015; Patzke et al. 2019; Beveridge et al. 2023). There is considerable evidence for both auxin and sugar models and no obvious reason that the two mechanisms should be mutually exclusive. Assuming that these two models may account for some, or all, of the apical dominance observed, determining their relative contributions should help inform and direct future research efforts.

The role of auxin in apical dominance has been tested across diverse approaches and species. Early studies found that removing the apical source of auxin promoted bud outgrowth, while supplying auxin to the cut stump could restore the suppression of branching (Thimann and Skoog, 1933). However, it was previously reported that supplemental auxin could not restore apical dominance in decapitated *Arabidopsis* (Cline, 1996). The current study revisited this issue and showed that apical dominance is effectively restored in *Arabidopsis* by applying auxin after decapitation. The disparate results are almost certainly due to the differing approaches used in the two studies. Here, decapitation and auxin supplementation were performed as soon as the cauline stem could be cut, before any discernable outgrowth of the rosette buds. Additionally, all cauline buds were removed and only the fate of the rosette buds was assessed. Finally, NAA in lanolin was used as the auxin source, which was previously shown to exert stronger effects than IAA (Cline, 1996). It is now apparent that auxin can suppress bud outgrowth in decapitated *Arabidopsis* as is the case for many other species. This conclusion aligns with other findings demonstrating an important role for auxin in *Arabidopsis* apical dominance. For instance, apically supplied auxin suppresses cauline bud growth in the split-plate assay used to assess auxin responsiveness (Chatfield et al. 2000). *Arabidopsis* mutants deficient in auxin signaling display greatly reduced apical dominance (Stirnberg et al. 1999), and auxin-overproducing lines show suppressed branching (Zhao et al. 2001). Similarly, manipulating auxin transport by rewiring PIN1 expression responses to auxin suppresses branching (Khakhar et al. 2018), while applying an auxin transport inhibitor to the cauline stem promotes the outgrowth of lower buds (Reddy at al., 2014). Modulating auxin responsiveness by altering the R:FR or manipulating phyB/PIF status has a major impact on rosette bud growth (Reddy et al. 2014; Holalu et al. 2020, 2021). Thus, there is abundant evidence that auxin is a major regulator of the *Arabidopsis* branching program.

The auxin transported basipetally in the PATS is expected to be sourced from the main shoot apex and from other lateral apices below it that feed auxin into the main stem PATS (Prusinkiewicz et al. 2009; Bennett et al. 2016). This collective auxin flow then suppresses the growth of inferior buds. If this model is correct, growth of the inferior rosette buds should be negatively correlated with the number of superior auxin-supplying cauline buds. Surprisingly, previous research suggested that rosette branching is not correlated with cauline branching (Fichtner et al. 2022). In contrast, removal of the cauline branches promoted the outgrowth of lower buds in the rosette, and intact plants demonstrated a highly significant negative correlation between cauline and rosette branching, as expected (Fig. 2). This relationship was robust to the unequal representation of cauline branch values in the dataset, as weighted least squares regression accounting for sample size differences across cauline branch classes yielded a comparable negative slope (Fig. S1). The discrepancy in the findings likely results from several factors. It should be noted that all cauline nodes produce branches. The number of cauline branches is thus determined by the number of cauline nodes, which in turn is correlated with flowering time (Hempel and Feldman, 1994; Fichtner et al. 2022). Because there is little variation in cauline node formation, a large dataset is required to capture a useful amount of variation. The association also appears to be sensitive to environmental parameters. The data used in the current study was generated by one laboratory, using similar, but not identical growth environments. Stronger correlations were generally observed within experimental sub-series conducted near the same time, likely reflecting small variations in growth practices or environments used by specific researchers. Plant health is extremely important when conducting physiological studies, especially in the case of branching. Extensive empirical data has shown that branching is suppressed when plant health is not optimal. The Col-0 accession routinely forms five or more branches when grown to a reasonable level of health in the environments typically employed in the laboratory. When Col-0 produces less branches, as is often reported, it likely reflects an unrecognized limitation or stress, making such results difficult to interpret. Our study also included only wild-type plants, as the hyperbranching mutants used to investigate branching display aberrant apical dominance, which logically defeats the possibility of determining correlation between cauline and rosette branching.

Although many lines of evidence support auxin regulation of apical dominance, the negative association between cauline and rosette branching may also be explained by the sugar branching model. One interpretation of this model (carbon-limited model) could be that sugars are diverted from supporting growth of lower rosette buds to promote the growth of cauline branches. Available sugars (carbon) would then limit the amount of growth from different parts of the plant. Another interpretation is that the sugars act not only as an energy and material source as in the carbon-limited model, but also as signals to modify the amount of rosette branching (sugar signaling model). Attributing the negative relationship between cauline and rosette branching to either sugar models and/or to a version of the classical PATS model is difficult, since perturbing one model’s parameters is likely to impact the other. Rather than address this issue directly, the current study instead sought to discover how sugar/carbon availability impacts *Arabidopsis* shoot architecture using the least complex and most easily interpreted systems possible.

We tested the effect of assimilate availability on Col-0 branching directly by removing all rosette leaves at the time of bolting, when the rosette buds were very small. The massive reduction in assimilate reserves and assimilation potential initially slowed bud growth but had essentially no impact on the final number of branches produced. Branch growth requires sugar-derived carbon for energy and materials, but the sugar status of the plant does not appear to be a strong determinant of the final branching morphology. Sufficient sugars necessary for branch development can apparently be provided by the growing bud/branch itself, thus obviating the need for plant-level sugar as an energy/material source or sugar status as a coordinating signal. These findings are consistent with the concepts of sectoriality and physiological carbon autonomy derived from research in both woody and herbaceous species, whereby developing branches neither import nor export carbon following an initial minor investment by orthostichous leaves (Sprugel et al. 1991; Marshall, 1996). Decapitation alone and in combination with defoliation promoted bud outgrowth over the corresponding treatments with intact cauline shoots suggesting that sugar-independent shoot-derived signals exert strong effects on branching (Fig. 6). Auxin is an obvious candidate for this signal.

Increasing PPFD promoted the shoot DW of all the accessions tested, but its effect on rosette bud outgrowth was weak in most accessions, indicating that branch numbers were determined to a greater extent by intrinsic programming rather than sugar supply. An earlier study of Col-0 found no difference in rosette branch numbers under low (160 μMoles m^-2^ s^-1^) or high (280 μMoles m^-2^ s^-1^) PPFD in soilless media, but a small increase in hydroponics (Su et al. 2011). Increasing PPFD had a more pronounced effect on the deposition of material into branches that formed and in some cases on the length of these branches than on the number of branches produced. The proportion of shoot mass allocated to rosette branches and buds was not changed across PPFD treatments, indicating that the additional assimilate was distributed among shoot components in the same proportions rather than redirected toward branching (Fig. S2e). Plants may prioritize the deposition of additional assimilate into a pre-programmed number of branches, in proportion to the growth of the shoot as a whole, over increasing the number of branches.

Van-0 differed from the other accessions in showing a moderately strong bud outgrowth response to increasing PPFD. In the path analysis, Van-0 was distinctive. Photoassimilate influenced branch number more than material deposition, the reverse of the pattern in the other accessions (Fig. 5b,c). Van-0 also had the shortest branches of the accessions tested and the most compact architecture (Fig. 5a). Using collection coordinates from the ABRC and climate data from WorldClim 2.1 (Fick and Hijmans, 2017), we found that Van-0 originates from a substantially wetter habitat than the other accessions, with an annual precipitation of ∼1700 mm, roughly 1.6- to 3.8-fold higher than that of the other accessions (442-1023 mm) (Fig. S8). It is tempting to speculate that in such an environment, allocating assimilate to additional branches rather than to reinforcing existing ones is favored, and that the short branches of Van-0 permit this by reducing the bending moment and thus the risk of mechanical failure.

Most of the accessions sampled flowered early under the long days used. These accessions displayed the typical basipetal pattern of bud activation described previously (Hempel and Feldman, 1994). The exception was Chi-0, which flowered relatively late. In this accession, lower buds often initiated during vegetative growth before the basipetal wave of bud initiation from the youngest nodes became evident, as documented previously in other lines (Grbic and Bleecker, 2000; Fichtner et al. 2022). Chi-0 consistently displayed the weakest associations between assimilation and branching parameters. The effect of carbon status on vegetatively initiated branches was not investigated separately, but there was no indication that carbon status was a positive determinant of branch initiation in late flowering plants. As in the early flowering lines, carbon status was positively associated with material deposition into Chi-0 branches, indicating the priority placed on this parameter in energetically favorable conditions.

These effects were tested in favorable conditions-possible that stresses/limitations could alter the outcome. Lower levels of light might produce different results, but we selected light levels that provided robust growth. Other studies have used relatively lower light levels, including in crops (Mason et al. 2014; Brewer et al. 2015; Nicolas et al. 2021; Fichtner et al. 2022).

Growing plants under varying PPFD permits the study of a largely unperturbed system, allowing the plant to reveal more naturally whether branch initiation is regulated by carbon availability.

Most accessions tell us that carbon is generally a minor regulator of the process.

## Methods and Materials

### Plant materials and growth conditions

Arabidopsis accessions including Bay-0 (CS57923), Chi-0 (CS1072), Col-0 (CS60000), Ob-0 (CS38905), Shadara (CS57924), and Van-0 (CS37007) were obtained from the ABRC. Seeds were stratified at 4°C for four days prior to sowing. Plants were grown in 180 mL round pots filled with synthetic medium composed of 2 parts peat moss, 1 part vermiculite and 1 part perlite adjusted to pH 5.7 with dolomitic lime. Pots were charged with modified Hoagland’s solution at sowing and then fertigated with modified Hoagland’s solution every day beginning 6 days after sowing, with increasing nutrient levels throughout the growth period. Plants were grown in growth chambers with 180 μMoles m^-2^ s^-1^ PPFD provided by t5 fluorescent lamps with a high R:FR ratio (>10) and an 18/6 h photo/thermoperiod with 24°C day/18°C night temperatures. Relative humidity was maintained at 60%. Plants that contributed historical data for the regression analysis of rosette branches vs cauline nodes were grown in a similar manner except that they were planted in 6 cell inserts (140 mL/cell) and were fertilized less intensively.

### Branch elongation analyses

Architectural characteristics and branch elongation of wild-type Col-0 were measured at 10 days post-anthesis (DPA) as described in Finlayson et al. (2010). The frequency of bud outgrowth was calculated as the proportion of buds at a given rosette position that grew to greater than 3 mm. Leaf number was counted, and the number of buds or meristems formed in each rosette axil was determined under a dissecting microscope. The lengths of the primary rosette and cauline branches (axis > 3 mm) and of the main inflorescence were measured. For each primary rosette and cauline branch, the number of nodes and the number of elongated secondary branches (axis > 3 mm) were recorded.

### Varying PPFD treatments

Varying PPFD treatments in a single growth chamber were obtained using baffles and neutral density filters. Pots were moved daily within light treatments to provide more even light exposure. Plants were initially grown with 180 μMoles m^-2^ s^-1^ PPFD for 10 days (or as specified) to establish a population of uniform plants. Plants were then matched by size and distributed to the different light treatments (120, 180 and 240 μMoles m^-2^ s^-1^ PPFD). At 10 days after anthesis plants were harvested and architectural parameters were measured using a ruler or digital calipers. Dissected plant parts were placed in foil envelopes and dried at 70°C to determine dry weight (DW). Specific branch mass and specific main stem mass were determined from the basal 50 mm segment of the longest rosette branch and of the main stem, respectively. Segments were dried and weighed as described above, and specific mass was expressed as dry weight per unit length (µg mm ¹).

### Decapitation and NAA treatment

The effect of auxin was assessed by decapitating plants at anthesis below the lowest cauline node, leaving about 10 mm of decapitated stem. Approximately 5 mg of lanolin containing 1% NAA was placed on the cut stem. Each day thereafter the NAA and a very thin slice (< 0.5 mm) of stem was removed and new auxin was reapplied. Mock treated plants received the same treatment except the lanolin did not contain NAA. Plants were harvested for analysis 10 days after anthesis.

### Removal of cauline branches, decapitation, and defoliation

The effect of cauline branch removal was assessed by removing all cauline buds at anthesis and then measuring architectural parameters at 10 days after anthesis. The effects of defoliation, decapitation and the combination of defoliation and decapitation, were assessed by decapitating the main shoot below the lowest cauline node, by removing all rosette leaves and by decapitating and defoliating when the main stem began to bolt. Architectural parameters were assessed at 12 days after anthesis (or estimated time of anthesis for plants that were decapitated).

### Path analysis

The relationships among PPFD, photoassimilate, and shoot branching architecture were modeled for each accession by piecewise structural equation modeling (SEM) using the piecewiseSEM package (v.2.3.1; Lefcheck, 2016) in R (v.4.5.2). Rosette leaf DW was used as a proxy for photoassimilate. The model specified PPFD as an effect on rosette leaf mass, and rosette leaf mass as an effect on three branching parameters: rosette branch number, total primary rosette branch length, and specific branch mass. Path coefficients were standardized to allow comparison across paths and accessions, and paths with *p* < 0.05 were considered significant.

### Climate data of accession origins

Reported collection coordinates for each accession were obtained from ABRC. Bioclimatic variables were extracted for these coordinates from WorldClim 2.1 at 2.5-arc-minute resolution (Fick and Hijmans, 2017) using the geodata and terra packages in R. Annual mean temperature (BIO1) and annual precipitation (BIO12) were retrieved for each site with terra::extract.

### Statistical analysis

Statistical analyses were performed in R using two tailed t-tests, one-way ANOVA, and linear regression. Where ANOVA indicated significant differences among treatments, means were separated using Tukey’s HSD test. Continuous data outliers were identified as data outside 1.5 times the interquartile range and were removed. To verify that results were not biased by unequal sample sizes across cauline branch classes, weighted least squares regression was also performed with weights set as the inverse of the within-class sample size. The regression data was normalized by dividing each parameter value by the sample mean which reveals relationships on a per plant basis but does not alter the coefficient of determination or the *p*-value. Significance was determined at α < 0.05.

## Author contributions

S.P. and S.A.F. designed the study. S.P. performed the research. S.P., S.A.F., and C.L. performed data analyses. S.P. wrote the manuscript with input from all authors.

## Acknowledgements

This work is dedicated to the memory of the late Dr. Scott A. Finlayson, with profound gratitude for his mentorship and unwavering support. This work was supported by Texas A&M AgriLife Research (S.A.F.). S.P. gratefully acknowledges Texas A&M AgriLife Research and the Department of Soil and Crop Sciences, Texas A&M University for their continued support following my graduation.

## References

Balla J, Kalousek P, Reinöhl V, Friml J, Procházka S (2011) Competitive canalization of PIN dependent auxin flow from axillary buds controls pea bud outgrowth. The Plant Journal 65(4): 571–577

Barbier F, Péron T, Lecerf M, Perez-Garcia M-D, Barrière Q, Rolčík J, Boutet-Mercey S, Citerne S, Lemoine R, Porcheron B, Roman H, Leduc N, Le Gourrierec J, Bertheloot J, Sakr S (2015) Sucrose is an early modulator of the key hormonal mechanisms controlling bud outgrowth in Rosa hybrida. Journal of Experimental Botany 66: 2569–2582

Barbier F, Dun EA, Kerr SC, Chabikwa TG, Beveridge CA (2019) An update on the signals controlling shoot branching. Trends in Plant Science 24(3): 220–236

Bennett T, Hines G, van Rongen M, Waldie T, Sawchuk MG, Scarpella E, Ljung K, Leyser O (2016) Connective Auxin Transport in the Shoot Facilitates Communication between Shoot Apices. PLoS Biology 14(4): e1002446

Beveridge CA, Rameau C, Wijerathna-Yapa A (2023) Lessons from a century of apical dominance research. Journal of Experimental Botany 74: 3903–3922

Brewer PB, Dun EA, Ferguson BJ, Rameau C, Beveridge CA (2009) Strigolactone Acts Downstream of Auxin to Regulate Bud Outgrowth in Pea and Arabidopsis. Plant Physiology 150: 482–493

Brewer PB, Dun EA, Gui R, Mason MG, Beveridge CA (2015) Strigolactone Inhibition of Branching Independent of Polar Auxin Transport Plant Physiology 168: 1820-1829

Chatfield SP, Stirnberg P, Forde BG, Leyser O (2000) The hormonal regulation of axillary bud growth in Arabidopsis. The Plant Journal 24: 159–169.

Cline MG (1996) Exogenous Auxin Effects on Lateral Bud Outgrowth in Decapitated Shoots. Annals of Botany 78: 255–266

Cline MG, Chatfield SP, Leyser O (2001) NAA Restores Apical Dominance in the axr3-1 Mutant of Arabidopsis thaliana. Annals of Botany 87: 61–65

de Jong M, George G, Ongaro V, Williamson L, Willetts B, Ljung K, McCulloch H, Leyser O (2014) Auxin and strigolactone signaling are required for modulation of Arabidopsis shoot branching by nitrogen supply. Plant Physiology 166(1): 384–95

Domagalska MA, Leyser O (2011) Signal integration in the control of shoot branching. Nature Reviews Molecular Cell Biology 12(4): 211–221

Fichtner F, Barbier FF, Feil R, Watanabe M, Annunziata MG, Chabikwa TG, Höfgen R, Stitt M, Beveridge CA, Lunn JE (2017) Trehalose 6 phosphate is involved in triggering axillary bud outgrowth in garden pea (Pisum sativum L.). The Plant Journal 92(4): 611–23

Fichtner F, Barbier FF, Annunziata MG, Feil R, Olas JJ, Mueller-Roeber B, Stitt M, Beveridge CA, Lunn JE (2021) Regulation of shoot branching in Arabidopsis by trehalose 6-phosphate. The New Phytologist 229(4): 2135–51

Fichtner F, Barbier FF, Kerr SC, Dudley C, Cubas P, Turnbull C, Brewer PB, Beveridge CA (2022) Plasticity of bud outgrowth varies at cauline and rosette nodes in Arabidopsis thaliana. Plant Physiology 188: 1586–1603

Fick SE, Hijmans RJ (2017) WorldClim 2: new 1 km spatial resolution climate surfaces for global land areas. International journal of climatology 37(12): 4302–4315

Finlayson SA, Krishnareddy SR, Kebrom TH, Casal JJ (2010) Phytochrome regulation of branching in Arabidopsis. Plant physiology 152(4): 1914–27

Grbic V, Bleecker AB (2000) Axillary meristem development in Arabidopsis thaliana. The Plant Journal 21: 215–223

Hempel FD, Feldman LJ (1994) Bi-directional inflorescence development in Arabidopsis thaliana: Acropetal initiation of flowers and basipetal initiation of paraclades. Planta 192: 276–286

Holalu SV, Reddy SK, Blackman BK, Finlayson SA (2020) PIF4 and PIF5 regulate axillary branching via bud abscisic acid and stem auxin signaling. Plant, Cell & Environment 43: 2224–2238

Holalu SV, Reddy SK, Finlayson SA (2021) Low Red light:Far Red light inhibits branching by promoting auxin signaling. J Plant Growth Reg 40: 2028–2036

Kebrom TH, Chandler PM, Swain SM, King RW, Richards RA, Spielmeyer W (2012) Inhibition of Tiller Bud Outgrowth in the tin Mutant of Wheat Is Associated with Precocious Internode Development. Plant Physiology 160: 308–318

Kebrom TH, Mullet JE (2015) Photosynthetic leaf area modulates tiller bud outgrowth in sorghum. Plant, Cell & Environment 38(8):1471–8

Khakhar A, Leydon AR, Lemmex AC, Klavins E, Nemhauser JL (2018) Synthetic hormone-responsive transcription factors can monitor and re-program plant development. eLife 7:e34702

Lefcheck JS (2016) piecewiseSEM: Piecewise structural equation modelling in r for ecology, evolution, and systematics. Methods in ecology and evolution 7(5):573–579

Mathan J, Bhattacharya J, Ranjan A (2016) Enhancing crop yield by optimizing plant developmental features. Development 143(18): 3283–3294

Marshall C (1996) Sectoriality and physiological organization in herbaceous plants: an overview. Vegetatio 127: 9–16

Mason MG, Ross JJ, Babst BA, Wienclaw BN, Beveridge CA (2014) Sugar demand, not auxin, is the initial regulator of apical dominance. Proceedings of the National Academy of Science 111: 6092–6097

McIntyre GI (1964) Mechanism of apical dominance in plants. Nature 203: 1190–1191

Nicolas M, Rodríguez-Buey ML, Franco-Zorrilla JM, Cubas P (2015) A recently evolved alternative splice site in the BRANCHED1a gene controls potato plant architecture. Current Biology 25(14): 1799–1809

Otori K, Tamoi M, Tanabe N, Shigeoka S (2017) Enhancements in sucrose biosynthesis capacity affect shoot branching in Arabidopsis. Bioscience, Biotechnology, and Biochemistry 81: 1470–1477

Otori K, Tanabe N, Tamoi M, Shigeoka S (2019) Sugar Transporter Protein 1 (STP1) contributes to regulation of the genes involved in shoot branching via carbon partitioning in Arabidopsis. Bioscience, Biotechnology, and Biochemistry 83: 472–481

Patil SB, Barbier FF, Zhao J, Zafar SA, Uzair M, Sun Y, Fang J, Perez-Garcia M-D, Bertheloot J, Sakr S, Fichtner F, Chabikwa TG, Yuan S, Beveridge CA, Li X (2022) Sucrose promotes D53 accumulation and tillering in rice. New Phytologist 234: 122–136

Patzke K, Prananingrum P, Klemens PAW, Trentmann O, Martins Rodrigues CM, Keller I, Fernie AR, Geigenberger P, Bolter B, Lehmann M, Schmitz-Esser S, Pommerrenig B, Haferkamp I, Ekkehard Neuhaus H (2019) The Plastidic Sugar Transporter pSuT Influences Flowering and Affects Cold Responses. Plant Physiology 179: 569–587

Prusinkiewicz P, Crawford S, Smith RS, Ljung K, Bennett T, Ongaro V, Leyser O (2009) Control of bud activation by an auxin transport switch. Proceedings of the National Academy of Sciences 106: 17431–17436

Reddy SK, Finlayson SA (2014) Phytochrome B promotes branching in Arabidopsis by suppressing auxin signaling. Plant Physiology 164: 1542–1550

Sprugel DG, Hinckley TM, Schaap W (1991) The theory and practice of branch autonomy. Annual Review of Ecology, Evolution, and Systematics 22: 309–334

Stirnberg P, Chatfield SP, Leyser HMO (1999) AXR1 acts after lateral bud formation to inhibit lateral bud growth in Arabidopsis. Plant Physiology 121: 839–847

Su H, Abernathy SD, White RH, Finlayson SA (2011) Photosynthetic photon flux density and phytochrome B interact to regulate branching in *Arabidopsis*. Plant, Cell & Environment 34: 1986–1998

Thimann KV, Skoog F (1933) Studies on the growth hormone of plants. III. The inhibitory action of the growth substance on bud development. Proceedings of the National Academy of Sciences 19: 714–716

Wang B, Smith SM, Li J (2018) Genetic regulation of shoot architecture. Annual Review of Plant Biology 69: 437–468

Zhao Y, Christensen SK, Fankhauser C, Cashman JR, Cohen JD, Weigel D, Chory J (2001) A role for flavin monooxygenase-like enzymes in auxin biosynthesis. Science 291: 306–309

