## Supplemental Figures for "Photosynthetic assimilate determines branch size and biomass more than branch number in Arabidopsis"

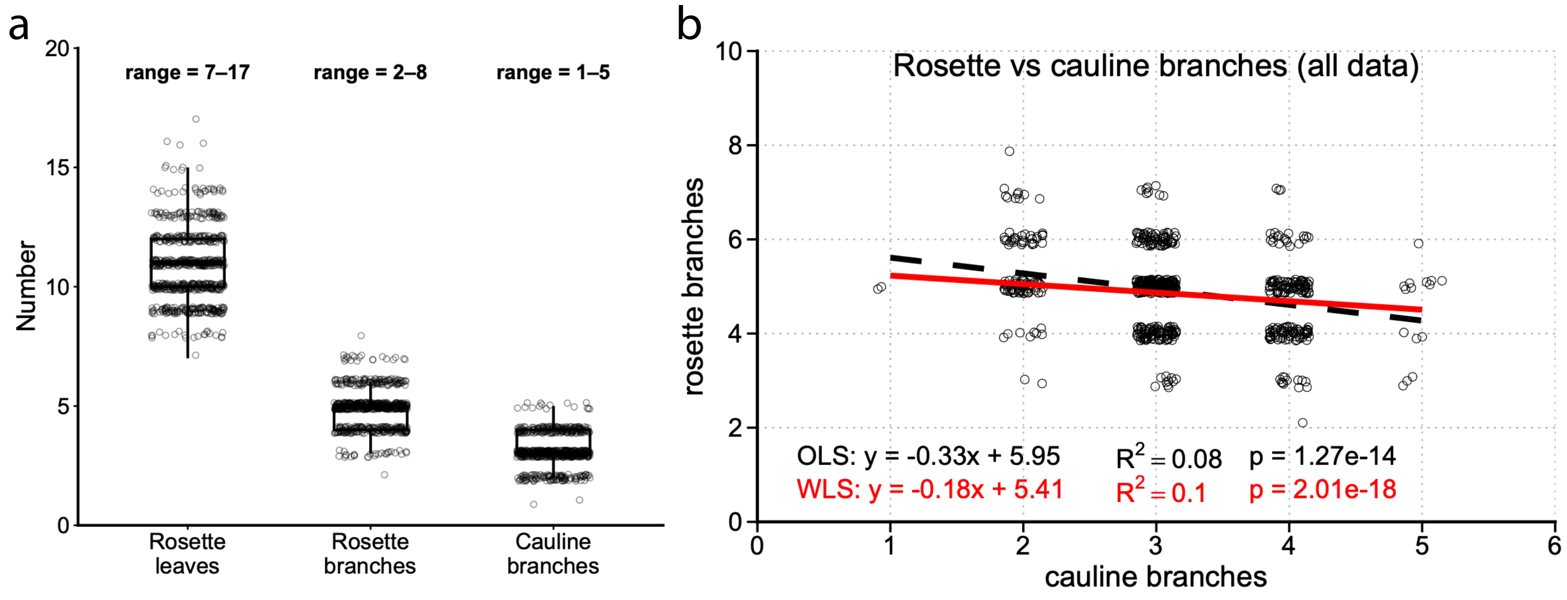


**Supplementary Figure 1.** Variation in rosette leaf, rosette branch, and cauline branch number in Col-0, and robustness of the cauline-rosette relationship to unequal sampling. **(a)** Distribution of rosette leaf numbers, rosette branch numbers, and cauline branch numbers across 718 Col-0 plants grown under closely matched conditions in 24 experiments. The observed range is given above each group. **(b)** Rosette branch numbers plotted against cauline branch numbers for the same dataset. The dashed black line is the ordinary least squares (OLS) regression and the solid red line the weighted least squares (WLS) regression, in which each observation was weighted by the inverse of the sample size of its cauline branch class. Regression equations, coefficients of determination, and *P*-values are given for each regression.

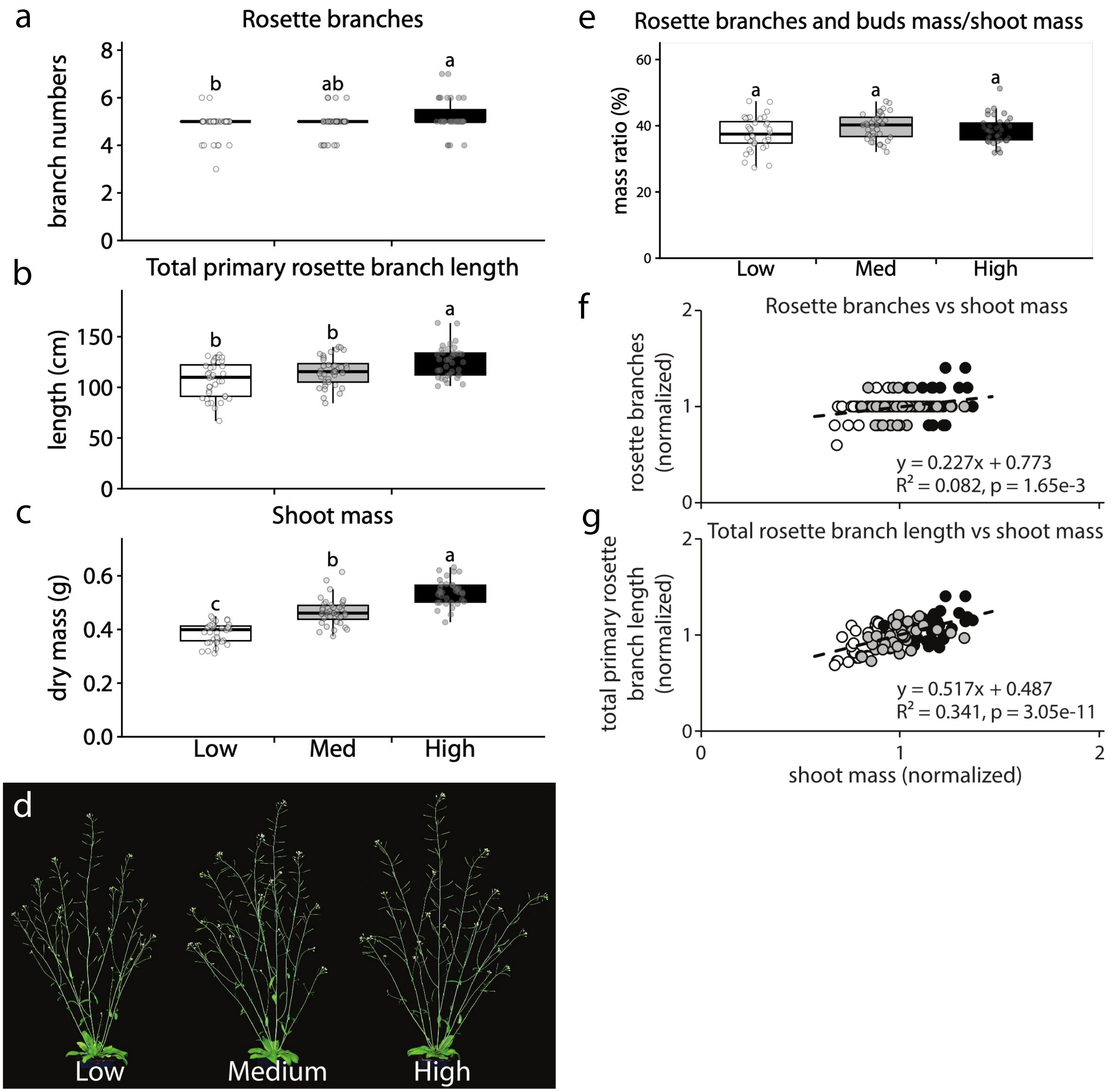


**Supplementary Figure 2.** The effects of photosynthetic assimilation on shoot architecture in Col-0. Plants are grown at low (120 μMoles $m^{-2}$ $s^{-1}$), medium (180 μMoles $m^{-2}$ $s^{-1}$), or high (240 μMoles $m^{-2}$ $s^{-1}$) PPFD from 20 days after sowing, the onset of bolting, and harvested 10 days after anthesis. **(a)** Rosette branch number. **(b)** Total primary rosette branch length. **(c)** Total shoot dry mass. **(d)** Representative Col-0 plants showing shoot architecture under varying PPFD treatment at harvest. **(e)** Rosette branch and bud dry mass expressed as a ratio of total shoot dry mass. **(f-g)** Regressions of rosette branch number (f) and total rosette branch length (g) against total shoot mass, each normalized to its sample mean. Each point represents a single plant, shaded by PPFD treatment: low (white), medium (grey), or high (black). Dashed lines are least-squares regressions, and the regression equation, coefficient of determination ($R^2$), and *P*-value are given for each regression. n = 36 plants per treatment. Different letters indicate significant differences between treatments (Tukey's HSD, $P < 0.05$).

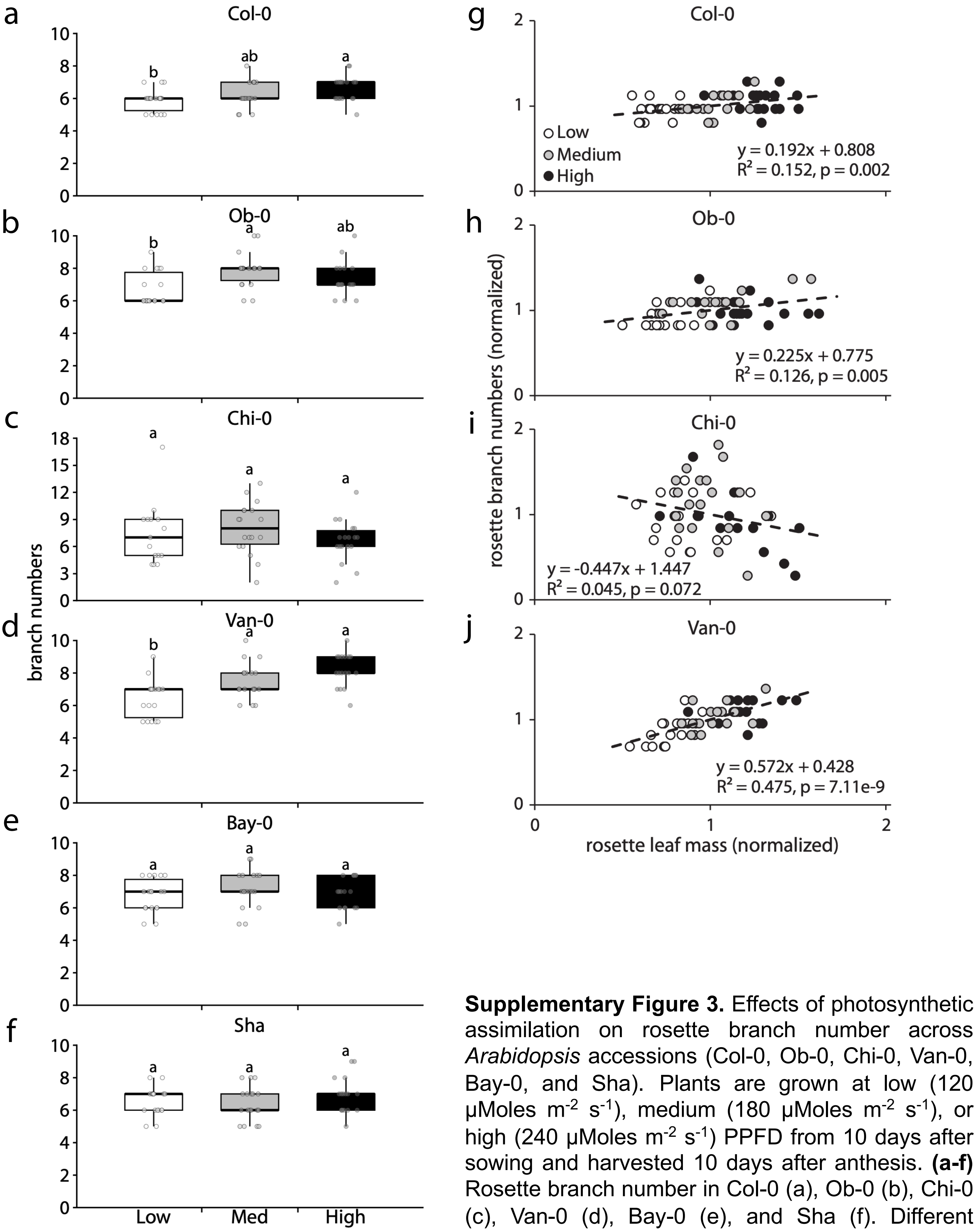


**Supplementary Figure 3.** Effects of photosynthetic assimilation on rosette branch number across *Arabidopsis* accessions (Col-0, Ob-0, Chi-0, Van-0, Bay-0, and Sha). Plants are grown at low (120 μMoles $m^{-2}$ $s^{-1}$), medium (180 μMoles $m^{-2}$ $s^{-1}$), or high (240 μMoles $m^{-2}$ $s^{-1}$) PPFD from 10 days after sowing and harvested 10 days after anthesis. **(a-f)** Rosette branch number in Col-0 (a), Ob-0 (b), Chi-0 (c), Van-0 (d), Bay-0 (e), and Sha (f). Different letters indicate significant differences between PPFD treatments (Tukey's HSD, $P < 0.05$). **(g-j)** Regressions of rosette branch number against rosette leaf mass, for Col-0 (g), Ob-0 (h), Chi-0 (i), and Van-0 (j), each normalized to its sample mean. Each point represents a single plant, shaded by PPFD treatment: low (white), medium (grey), or high (black). Dashed lines are least-squares regressions, and the regression equation, coefficient of determination ($R^2$), and *P*-value are given for each regression. n = 51-54 plants per accession (n = 17-18 per PPFD treatment).

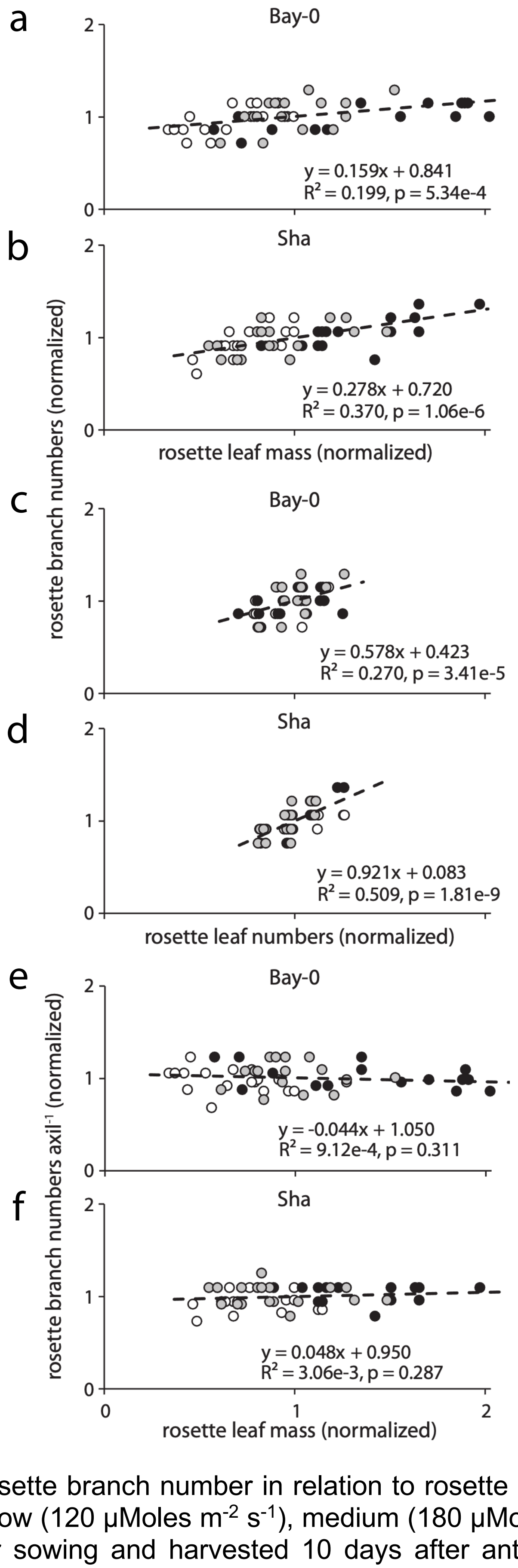


**Supplementary Figure 4.** Rosette branch number in relation to rosette leaf mass and leaf number in Bay-0 and Sha. Plants are grown at low (120 µMoles $m^{-2}$ $s^{-1}$), medium (180 µMoles $m^{-2}$ $s^{-1}$), or high (240 µMoles $m^{-2}$ $s^{-1}$) PPFD from 10 days after sowing and harvested 10 days after anthesis. **(a-b)** Regressions of rosette branch number against rosette leaf mass for Bay-0 (a) and Sha (b). **(c-d)** Regressions of rosette branch number against rosette leaf number for Bay-0 (c) and Sha (d). **(e-f)** Regressions of rosette branch number per axil against rosette leaf mass for Bay-0 (e) and Sha (f). All parameters were normalized to their sample means. Points represent individual plants: low (white), medium (grey), or high (black) PPFD. Dashed lines are least-squares regressions, and the regression equation, coefficient of determination ($R^2$), and *P*-value are given for each regression. n = 54 plants per accession (n = 18 per PPFD treatment).

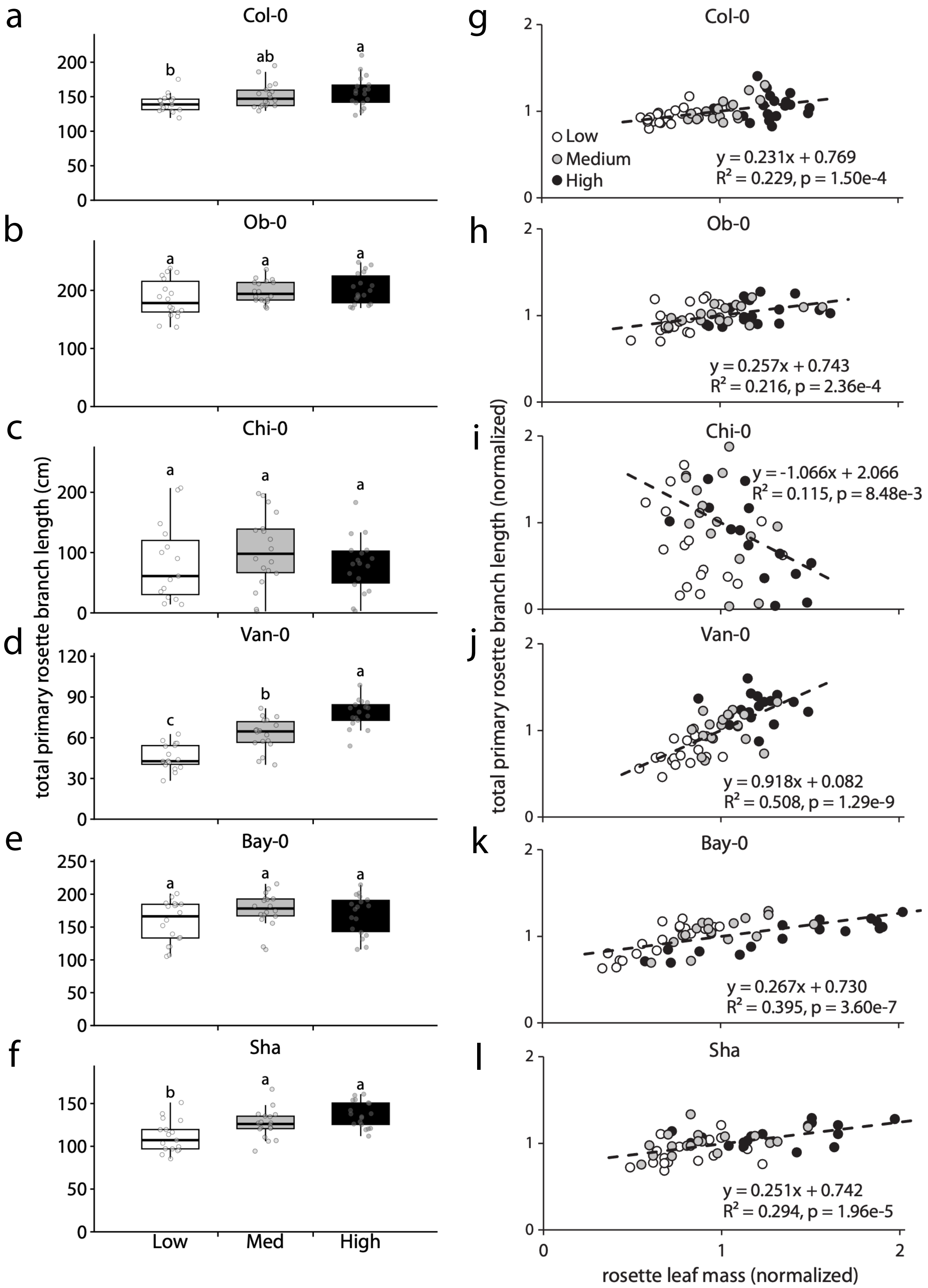


**Supplementary Figure 5.** Effects of photosynthetic assimilation on total primary rosette branch length across *Arabidopsis* accessions (Col-0, Ob-0, Chi-0, Van-0, Bay-0, and Sha). Plants are grown at low (120 µMoles $m^{-2}$ $s^{-1}$), medium (180 µMoles $m^{-2}$ $s^{-1}$), or high (240 µMoles $m^{-2}$ $s^{-1}$) PPFD from 10 days after sowing and harvested 10 days after anthesis. **(a-f)** Total primary rosette branch length in Col-0 (a), Ob-0 (b), Chi-0 (c), Van-0 (d), Bay-0 (e), and Sha (f). Different letters indicate significant differences between PPFD treatments (Tukey's HSD, $P < 0.05$). **(g-l)** Regressions of total primary rosette branch length against rosette leaf mass, for Col-0 (g), Ob-0 (h), Chi-0 (i), and Van-0 (j), Bay-0 (k), and Sha (l), each normalized to its sample mean. Each point represents a single plant, shaded by PPFD treatment: low (white), medium (grey), or high (black). Dashed lines are least-squares regressions, and the regression equation, coefficient of determination ($R^2$), and $P$-value are given for each regression. n = 51-54 plants per accession (n = 17-18 per PPFD treatment).

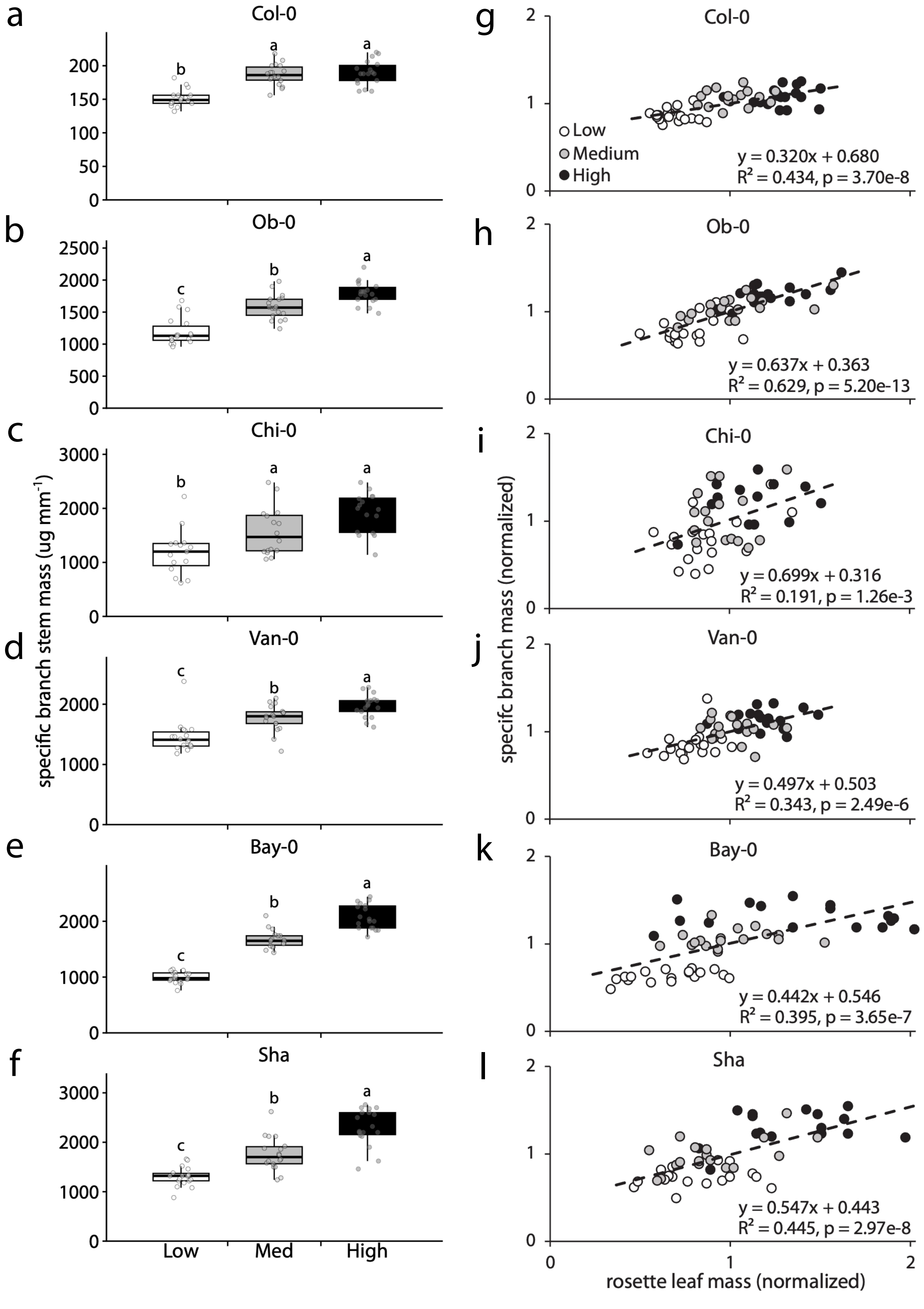


**Supplementary Figure 6.** Effects of photosynthetic assimilation on specific branch mass across *Arabidopsis* accessions (Col-0, Ob-0, Chi-0, Van-0, Bay-0, and Sha). Plants are grown at low (120 μMoles $m^{-2}$ $s^{-1}$), medium (180 μMoles $m^{-2}$ $s^{-1}$), or high (240 μMoles $m^{-2}$ $s^{-1}$) PPFD from 10 days after sowing and harvested 10 days after anthesis. **(a-f)** Specific branch mass in Col-0 (a), Ob-0 (b), Chi-0 (c), Van-0 (d), Bay-0 (e), and Sha (f). Different letters indicate significant differences between PPFD treatments (Tukey's HSD, $P < 0.05$). **(g-l)** Regressions of specific branch mass against rosette leaf mass, for Col-0 (g), Ob-0 (h), Chi-0 (i), and Van-0 (j), Bay-0 (k), and Sha (l), each normalized to its sample mean. Each point represents a single plant, shaded by PPFD treatment: low (white), medium (grey), or high (black). Dashed lines are least-squares regressions, and the regression equation, coefficient of determination ($R^2$), and $P$-value are given for each regression. n = 51-54 plants per accession (n = 17-18 per PPFD treatment).

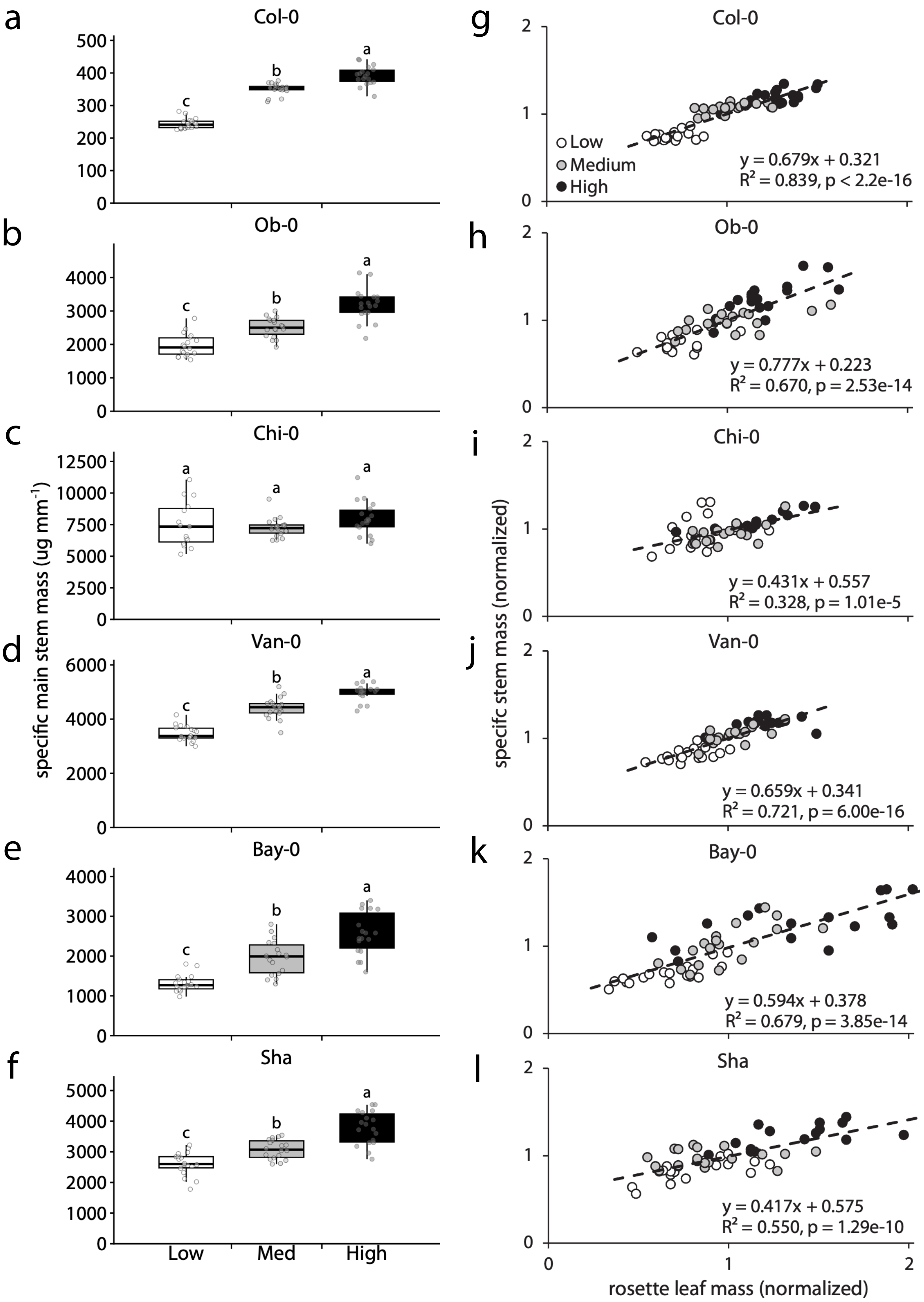


**Supplementary Figure 7.** Effects of photosynthetic assimilation on specific main stem mass across *Arabidopsis* accessions (Col-0, Ob-0, Chi-0, Van-0, Bay-0, and Sha). Plants are grown at low (120 µMoles $m^{-2}$ $s^{-1}$), medium (180 µMoles $m^{-2}$ $s^{-1}$), or high (240 µMoles $m^{-2}$ $s^{-1}$) PPFD from 10 days after sowing and harvested 10 days after anthesis. **(a-f)** Specific main stem mass in Col-0 (a), Ob-0 (b), Chi-0 (c), Van-0 (d), Bay-0 (e), and Sha (f). Different letters indicate significant differences between PPFD treatments (Tukey's HSD, $P < 0.05$). **(g-l)** Regressions of specific main stem mass against rosette leaf mass, for Col-0 (g), Ob-0 (h), Chi-0 (i), and Van-0 (j), Bay-0 (k), and Sha (l), each normalized to its sample mean. Each point represents a single plant, shaded by PPFD treatment: low (white), medium (grey), or high (black). Dashed lines are least-squares regressions, and the regression equation, coefficient of determination ($R^2$), and *P*-value are given for each regression. n = 51-54 plants per accession (n = 17-18 per PPFD treatment).

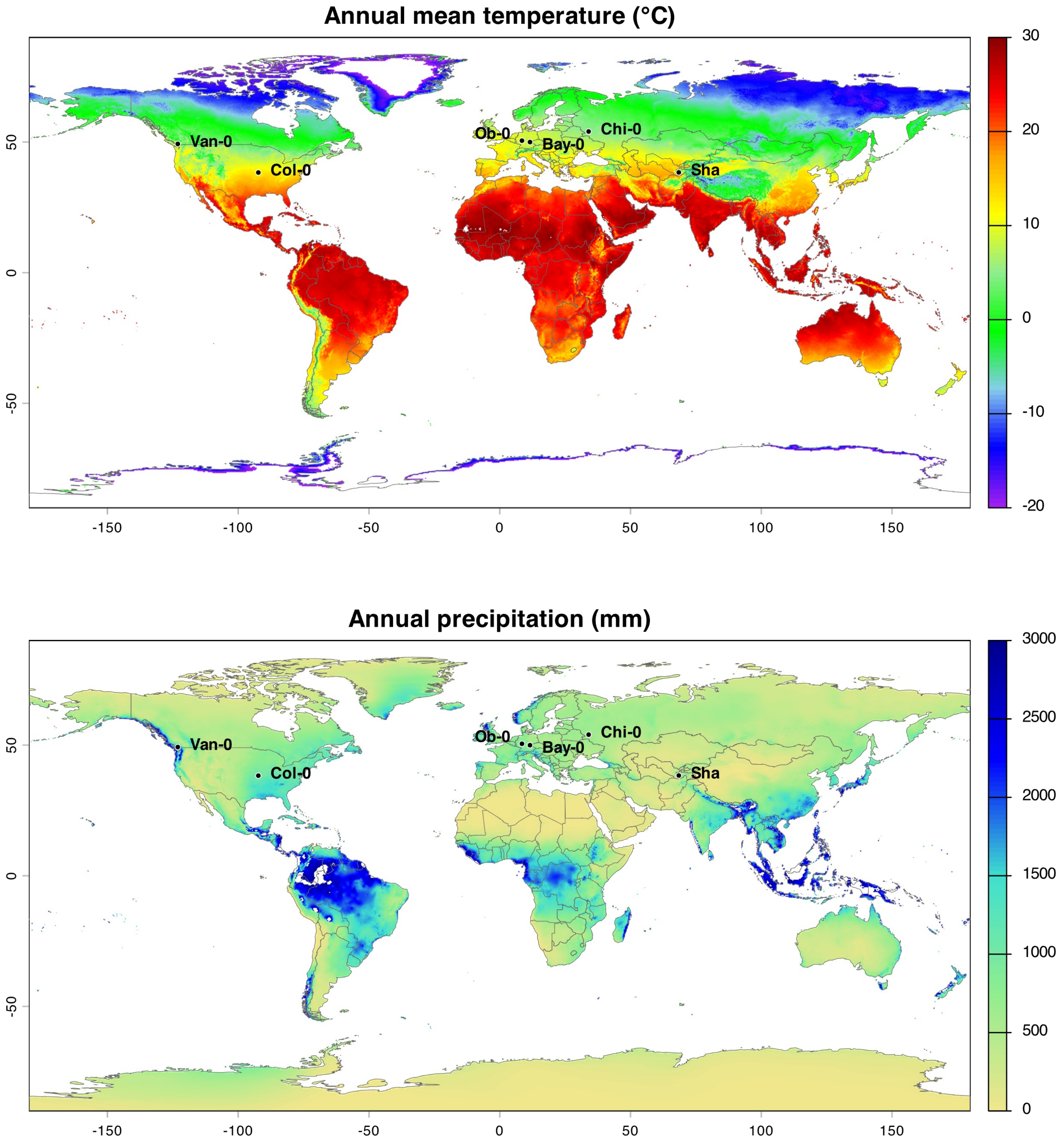


**Supplementary Figure 8.** Climatic context of the accessions used in this study. Annual mean temperature (upper panel) and annual precipitation (lower panel) from WorldClim 2.1 at 2.5-arc-minute resolution. Points mark the reported collection coordinates of Col-0, Ob-0, Chi-0, Van-0, Bay-0, and Sha, obtained from Arabidopsis Biological Resource Center (ABRC).
